# Directed differentiation of Dental Pulp Pluripotent-like Stem Cells into Hepatocyte-like Cells

**DOI:** 10.1101/2020.12.09.418780

**Authors:** Carlos Gil-Recio, Sheyla Montori, M Cámara Vallejo, Saddam Al Demour, Eduard Ferrés-Padró, Miguel Barajas, Carles Martin, Ashraf Al Madhoun, Maher Atari

**Affiliations:** Regenerative Medicine Research Institute, UIC Barcelona, Barcelona, Spain; Oral and Maxillofacial Surgery Department, Hospital Clinico de Barcelona, Barcelona, Spain; Department of Special Surgery/Division of Urology, The University of Jordan, School of Medicine, Amman, Jordan; Oral and Maxillofacial Surgery Department, Fundació Hospital de Nens de Barcelona, Barcelona, Spain; Biochemistry and Molecular Biology Department, Universidad Pública de Navarra, Pamplona, Spain; Functional Genomic Unit, Dasman Diabetes Institute, Kuwait; Surgery and Oral Implantology Department, College of Dentistry Universitat Internacional de Catalunya, Spain

**Keywords:** hepatocyte differentiation, adult stem cells, dental pulp pluripotent stem cells, DPPSCs

## Abstract

Liver diseases are a major cause of morbidity and mortality. Dental pulp pluripotent-like stem cells (DPPSC) are a considerable promise in tissue engineering and regenerative medicine as a source of tissue-specific cells, therefore this study aims to demonstrate their ability to generate functional hepatocyte-like cells *in vitro*. Cells were differentiated on a collagen scaffolds in serum free media supplemented with growth factors and cytokines to recapitulate liver development. At day 5, the differentiated DPPSC cells expressed the endodermal markers FOXA1 and FOXA2. Then, the cells were derived into the hepatic lineage generating hepatocyte-like cells. In addition to the associated morphological changes, the cells expressed the hepatic genes HNF6, AFP. The terminally differentiated hepatocyte-like cells expressed the liver functional proteins albumin and CYP3A4. In this study, we report an efficient serum-free protocol to differentiate DPPSCs into functional hepatocyte-like cells. Our approach promotes the use of DPPSCs as a new source of adult stem cells for prospective use in liver regenerative medicine.

## 1 Introduction

The liver is the largest internal organ providing essential metabolic, exocrine, and endocrine metabolites. Hepatocytes and cholangiocytes, the bile duct epithelial cells, are parenchymal cells which comprise approximately 70% of the adult liver mass. Both cell types are derived from the embryonic definitive endoderm [1]. The liver is the main body homeostasis regulator; therefore, liver diseases cause high morbidity and mortality rates. The incidence of end-stage liver disease (ESLD) is increasing worldwide [2] due to alcohol induced liver disease, nonalcoholic fatty liver disease, or hepatitis. Orthotopic liver transplantation (OLT) is the current optimal treatment with more than 70% of the overall 5-year survival rate for certain ESLD cases [3]. OLT obstacles include donor availability, surgical risk, high costs, and the requirement for life-long immunosuppressors [4; 5; 6]. Alternatively, hepatocyte transplantation (HT) is a promising procedure, less invasive, can be performed repeatedly, used mainly to replace an inborn deficient metabolic function [7] [8]. Yet, HT hurtles include donor availability, low cell engraftment, and longevity [9]. Therefore, the scientific community has put an effort to improve cell-based hepatocyte generation.

Dental pulp stem cells (DPSCs) are a multipotent heterogeneous cell population embedded within the pulp cavity of impacted third molars. DPSCs were initially isolated and characterized by Gronthos *et. al*. [10]. Subsequently, several investigators reported DPSCs’ isolation, characterization, differentiation, and banking [11; 12]. In comparison with embryonic and other adult stem cells, DPSCs are obtained from disposable dental pulp after occlusion management, which is considered as a medical waste, noninvasive isolation procedure, not associated with ethical constraints, and no risk of teratoma formation. Furthermore, DPSCs do not compromise their stemness, viability, proliferation, or differentiating capabilities after cryopreservation [13; 14]. Therefore, DPSCs have the potential to be a promising personalized patient-specific stem cell source for regenerative therapy. Nevertheless, a major concern is that DPSCs comprise progenitor cell populations that are marked by diverse characteristics, such as clonal heterogeneity, multilineage differentiation, self-renewal capacity and phenotypic complexity.

In the context of cell-based hepatogenesis, DPSCs-mediated hepatocyte-like generations has advanced rapidly because of the previously accumulated differentiation protocols applied for human embryonic stem cells (hESCs) and bone marrow mesenchymal cells (BM-MSCs) as prospective sources for regenerative hepatocytes [15; 16; 17; 18; 19; 20]. Implementing defined, serum-free and stepwise differentiation protocols that mimic hepatocytes’ developmental stages during embryogenesis are sufficient for inducing hepatogenesis. Using this approach, Ishkitiev *et. al*. were the pioneers to demonstrate the hepatogenic differentiation potential of DPSCs. Initially, Ishkitiev *et. al*. developed two staged conditioned media containing a low percentage of fetal bovine serum (FBS) but enriched with essential hepatogenic inducers [21]. Later, they used serum-free conditioned media to generate hepatocyte-like cells from CD117^+^ DPSCs subpopulation [22]. The latter hepatogenesis protocol utilized three developmental stages, i.e., cell specification, differentiation, and maturation, to generate cells with morphological, phenotypical, and functional characteristics like hepatocytes. This approach was further improved and implemented in later studies [11; 23]. Nevertheless, the Heterogeneity nature of the DPSCs influences their differentiation capacity and yield, therefore, we focused our efforts to identify and characterize the differentiation potential of a defined subpopulation of DPSCs with pluripotent-like phenotype [24; 25; 26; 27; 28].

Dental pulp pluripotent stem cells (DPPSCs) are a unique sub-population of DPSCs [25], expressing the pluripotency markers Oct4A, NANOG, and SOX2 [24; 29; 30]. DPPSCs exhibits a normal human karyotype with no aneuploidy, polyploidy, or any chromosomal abnormality during the metaphase stage even after more than 65 passages [24; 31]. Unlike other adult stem cells, DPPSCs form teratoma-like structures when injected subcutaneously in immunodeficient mice, and generate embryonic bodies like in *in vitro* cultures, which are exclusive phenotypes of hESCs and induced pluripotent stem cells (iPSCs) iPSCs [24; 32; 33; 34]. Furthermore, DPPSCs can be differentiated into cells from each of the three germ-embryonic layers, including osteocytes and bone- [25; 26], endothelial- and neural-like cells [25; 26], smooth and skeletal muscles [27].

In this study, we evaluated the adult DPPSCs-mediated hepatogenic capacity using a novel serum free-directed-step wide differentiation protocol and collagen scaffolds to induce 3-dimentaial (3D) culture conditions. DPPSCs were first differentiated into definitive endoderm (DE), which were then directed to the hepatogenic lineage. We observed that treatment of DPPSCs, seeded on scaffolds with Activin A (Act A), Wnt Family Member 3A (Wnt3A), and knockout serum replacement (KOSR) conditioned media, enhances their commitment capacity toward DE- lineage. Subsequently, treatment with fibroblast growth factor 4 (FGF4) and hepatocyte growth factor (HGF) induced gut tube formation, early hepatogenic markers, and finally, functional hepatocyte-like cells were generated in conditional differentiation medium supplemented with HGF, FGF4, dexamethasone (Dex) and oncostatin M (OSM). Preliminary results were previously published in a preprint [35].

## 2. Materials and methods

### 2.1. Ethics Approval and Statement

All research in the present study was conducted in accordance with the code of ethics of the World Medical Association for experiments involving humans (Helsinki Declaration of 1975) and the guidelines on human stem cell research issued by the Committee of Bioethics at the Universitat International de Catalunya, Spain.

### 2.2. DPPSCs Culture and Maintenance

DPPSCs clones were previously isolated and characterized by our group as previously described [24; 31]. In Brief, immediately after extraction, the third molars were vigorously washed with 70% ethanol and sterile distilled water. The molar pulp tissues were extracted, and cells were isolated by digesting the pulp tissue with collagenase type I (3 mg/ml, Sigma-Aldrich, Germany) for 60 minutes at 37°C. Then, cells were separated mechanically with an insulin syringe and centrifuged for 10 minutes at 1800 rpm. Primary cell lines were established, the medium was changed every 4 days, and cell density was maintained at a low density of 80-100 cells/cm^2^. DPPSCs were cultured in precoated flasks with 100 ng/ml fibronectin (Life Technologies, Waltham, MA, USA) in a medium consisting of 60% DMEM-low glucose (Life Technologies) and 40% MCDB-201 (Sigma-Aldrich); supplemented with 1X SITE Liquid Media Supplement (Sigma-Aldrich); 1X linoleic acid-bovine serum albumin (LA-BSA, Sigma-Aldrich); 10^−4^ M L-ascorbic acid 2-phosphate (Sigma-Aldrich); 1X Penicillin-Streptomycin (Life Technologies); 2% fetal bovine serum (FBS, Sigma-Aldrich); 10 ng/ml hPDGF-BB (Abcam, Waltham, MA, USA); 10 ng/ml EGF (Sigma-Aldrich); 1000 U/ml LIF (Millipore, USA); chemically defined Lipid Concentrate (Life Technologies); 0.8 mg/ml BSA (Sigma-Aldrich) and 55 μM β-mercaptoethanol (Sigma-Aldrich).

### 2.3. Definitive Endoderm Induction using Growth Factors

For the DE- induction, 5×10^4^ cells/cm^2^ were seeded on 6-well plates (Cell Coat, UK) and cultured in RPMI medium (Mediatech, Los Altos, CA, USA) containing GlutaMAX (Life Technologies), penicillin/streptomycin, 0.5% defined fetal bovine serum (FBS, HyClone, Cytiva Europe GmbH, Spain) and supplemented with each one of the following combinations of growth factors: (i) 100 ng/ml Act A (R&D Systems, UK), (ii) 100 ng/ml Act A + 50 ng/ml Wnt3A (R&D Systems), (iii) 100 ng/ml Act A + 50 ng/ml bone morphogenetic protein-4 (BMP4, R&D Systems), (iv) 100 ng/ml Act A + 50 ng/ml Wnt3A + 50 ng/ml BMP4, (v) 100 ng/ml Act A + 10 ng/ml FGF4 (R&D Systems) or (vi) 100 ng/ml Act A + 10 ng/ml basic-FGF (bFGF, Life Technologies). Three days post-induction, the medium was changed to the same RPMI-based medium and supplements except for the replacement of FBS with 2% KOSR (Life Technologies) and the incubation was maintained for another 2 days.

### 2.4. Definitive Endoderm Induction Using Different Biomaterials

Application of biomaterials to cell culture has been shown to enhance cellular programming and differentiation. In this study, we used collagen I or fibronectin, the most abundant components of the extracellular matrix that provide a structural network for the cells microenvironment and were reported to enhance DE- differentiation *in vitro* [36; 37]. Using the sample induction media described above, cells were seeded on plates coated with collagen type I, 100 ng/ml fibronectin, or were co-cultured with HepG2 cells (20,000 cell/cm^2^), in addition to the controls.

### 2.5. Hepatogenic differentiation

After setting the optimal conditions for DE-induction; we aimed to generate stepwise functional hepatocyte-like cell. The gut tube and hepatocyte specifications were initiated using RPMI medium supplemented with 2% KOSR, 10 ng/ml FGF4, and 10 ng/ml HGF (R&D Systems). Three days later, RPMI was replaced with the enriched minimum essential medium (MEM, Sigma-Aldrich) supplemented with 1% BSA, 10 ng/ml FGF4, and 10 ng/ml HGF. Three days later, hepatocyte maturation was induced using complete hepatocyte culture medium (HCM, Single-quote, Lonza, USA) with all supplements according to the manufacturer recommendations; containing 10 ng/ml FGF4, 10 ng/ml HGF, 10 ng/ml OSM and 10^−7^ M Dex (Sigma-Aldrich). The hepatocyte maturation was proceeded for nine days as previously reported by Ishkitiev *et. al*. [21; 22]. The differentiation protocol was performed for a total of 22 days, and the media were refreshed every 2-3 days.

### 2.6. Immunofluorescence microscope imaging

Immunofluorescence was performed as described previously [38; 39]. Briefly, cells were cultured overnight on glass coverslips coated with collagen I, fixed with 4% paraformaldehyde for 15 min, and permeabilized with 10% Triton X-100 for 30 min. Then, cells were blocked with 5% BSA in PBS for 30 min and incubated for one hour with the primary antibodies: alpha-1 antitrypsin (AAT; Abcam) or albumin (Alb; Abcam). Followed by one-hour incubation with prospective secondary antibodies (Abcam). Between each step, cells were washed with 1% BSA in PBS. Fluorescent images were captured using a fluorescence microscopy Olympus AX70 (Olympus Optical, Tokyo, Japan) as previously described [40; 41].

### 2.7. Immunocytochemistry

Cells were fixed with Thin Prep-CytoLyt solution for 30 min at room temperature and centrifuged at 5,000 rpm for 5 min. Cell’s pellet was solubilized in Thin prep-CytoLyt solution and then resuspended in PreservCyt solution (Thinprep) for 15 min and processed on a ThinPrep 2000 processor. The microscope slides harboring the processed cells were fixed with 96% ethanol, washed twice with distilled water, and blocked in 0.5% hydrogen peroxide/methanol for 10 min. Immunostainings were performed using a Leica Bond-MAX automated IHC and ISH *Stainer* in accordance with the manufacturer’s instructions. Briefly, the slides were washed with bond wash solution (Biosystems, USA) and the antigens retrieval procedure was performed in accordance with Bond heat standard protocol (ER1) using citric buffer, pH 6, for 30 min at 95°C. Slides are washed and treated with post-primary solution for 8 min. Next, the slides were incubated in Polymer AP for 8 min and washed. Then, the mixed Diaminobenzidine Refine solution was applied for 10 min. Samples were counterstained with water hematoxylin for 5 min. Non-immune immunoglobulins of the same isotype as the primary antibodies were used as a control for each experiment.

### 2.8. RNA Extraction, cDNA Synthesis and qRT-PCR Reactions

Total RNA was extracted from cells using the total RNA purification Trizol Reagent (Life Technologies) in accordance with the manufacturer’s protocol. RNA was quantified using a NanoDrop 2000c spectrophotometer (Thermo Fisher Scientific, Waltham, MA, USA) and RNA integrity was evaluated by 2% agarose gel electrophoresis [42]. First strand cDNA was synthesized from 200 ng RNA by reverse transcription using QuantiTect Reverse Transcription Kit (Qiagen Inc., Germantown, MD, USA). qRT-PCR reactions were performed as previously described [43; 44]. Primer pairs with equivalent efficiencies (Table 1) were selected from Primer Bank [45] or designed using primer-BLAST tools (http://www.ncbi.nlm.nih.gov/tools/primer-blast/) [46]. qRT-PCR was performed on the ABI7900 system (Applied Biosystems, USA) using SDS software. Relative gene expression was calculated using a comparative Ct method as previously described [47; 48]. Results were normalized to GAPDH CT-values, and the relative expressions were determined relative to control or Day 0 undifferentiated cells. Data show Means ± standard deviation. Total liver RNA (Life Technologies) was used as a positive control.

**Table 1.**
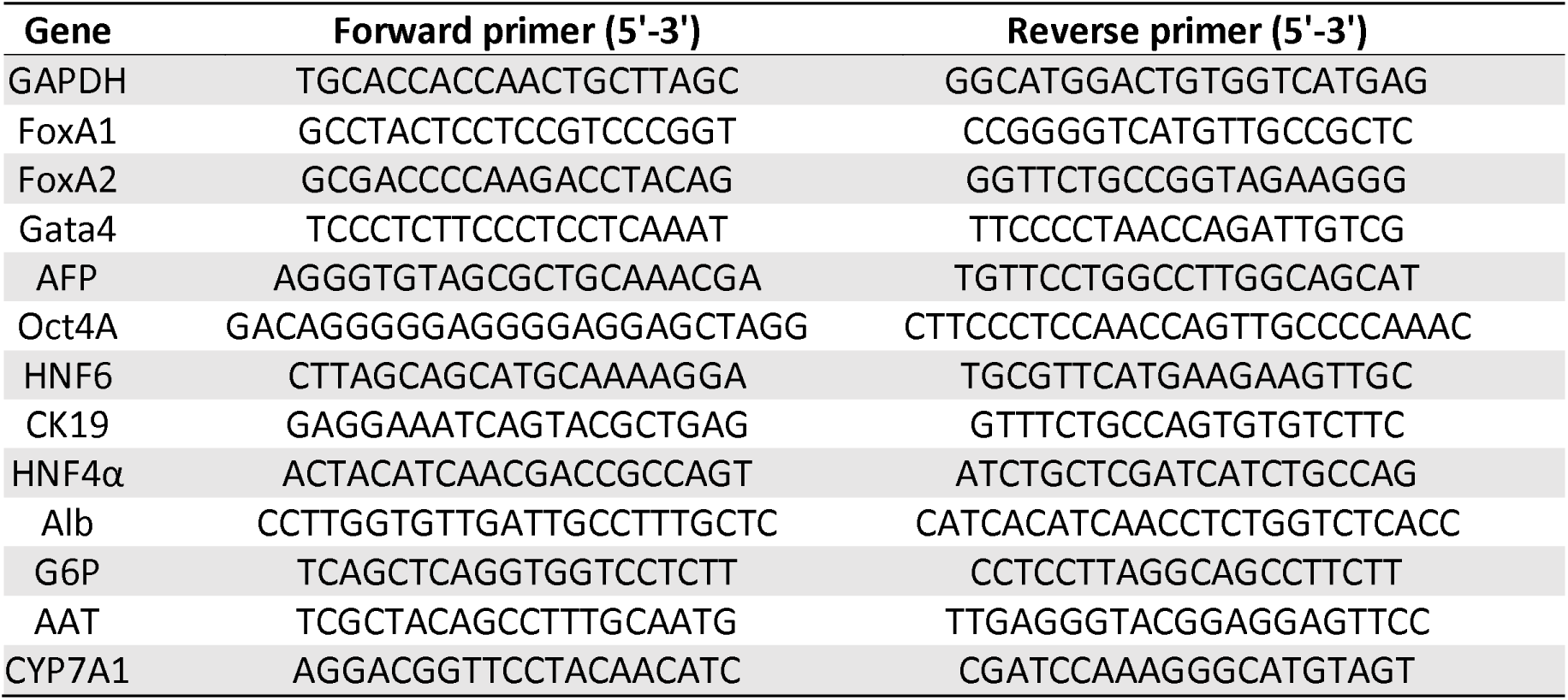
Oligonucleotide sequences of primers utilized for real-time qRT-PCR.

### 2.9. Hepatic Biochemical Analysis of Supernatants

Hepatic enzymatic profiles for aspartate transaminase (AST), alanine aminotransferase (ALT), alkaline phosphatase (ALP) and gamma-glutamyl transferase (GGT) were analyzed from supernatant extracts of the different samples. The activity was measured by specific colorimetric detection kit (Linear Chemicals, Spain) at 25°C in accordance with the manufacturer’s instructions. The specific compounds used for the following kinetics of the reaction were as the following: for AST and ALT, the oxidation of NADH was measured at light *absorbance* at 340 nm; for ALP, the formation of 4-Nitrophenol was measured at light *absorbance* at 405 nm; for GGT, the catalysis of γ-glutamyl-3-carboxy-4-nitroanilide was measured at light *absorbance* at 410 nm using spectrophotometer (Thermo Fisher Scientific).

### 2.10. Cytochrome P450 3A4 Metabolic Activity Assay

Cytochrome P450 3A4 (CYP3A4) enzyme activity assays were assessed by measurement of luciferase activity using P450-Glo CYP3A4 assay kit (Promega, Madison, WI, USA) in accordance with the manufacturer’s instructions. Differentiated and undifferentiated control DPPSC cells were treated with 25 μM Rifampicin for 48 hrs. Then, treated cells were incubated with a fresh serum-free medium containing 50 μM Luciferin PFBE with or without 5 μM Erythromycin (an antagonist) for 30 min at 37°C. Then, 50 μL of the medium was transferred into a 96-well plate and mixed with 50 μL luciferin detection reagent to initiate the luminescent reaction. After 20 min incubation at room temperature, the luminescence was measured with a luminometer (biotek, USA).

### 2.11. Albumin Assay

Alb secretion was measured by using an Albumin Fluorescence Assay Kit (Fluka, USA) in accordance with the manufacturer’s instructions. Briefly, a calibration curve was generated using the kit- supplied standard human Alb concentrations. These calibration samples were mixed with albumin Blue 580 reagent pre-diluted in buffer solution. The fluorescence was measured in a spectrophotometer (*absorbance* wavelength peal, λex = 600 nm, and λem = 630 nm. Differentiated and undifferentiated control DPPSCs were mixed with the same reagents and the fluorescence was measured under the same conditions. Results are extrapolated to the calibration curve. Conditioned medium was collected from equivalent numbers of cells.

### 2.12. Periodic acid–Schiff staining for glycogen accumulation

At differentiation day 22, differentiated and undifferentiated control DPPSCs were fixed in 4% formaldehyde for 15 min at room temperature. After two washing steps with PBS, cells were incubated in 1% periodic acid for 5 min, then washed with distilled water. The treated cells were incubated with Schiff’s reagent (Sigma-Aldrich) for 15 min. After a 10 min wash in distilled water, hematoxylin counterstain solution was applied for 90 sec and washed with distilled water.

### 2.13. Statistical analysis

Results are reported as mean ± standard deviation (see Figure legend for specific details regarding the number of biological replicates, independent experiments, and technical replicates). Statistics were performed using the Statgraphics XVI software. The methods used were two-tailed Student’s t-test and ANOVA for multiple factors. Values with p<0.05 were considered statistically significant.

## 3. Results

### 3.1. Definitive Endoderm Induction

Using hESCs, iPSCs, and human umbilical cord Wharton’s jelly mesenchymal stem cells, studies have shown that Act A is the main DE induction factor [49; 50; 51; 52]. Therefore, we tested this concept on DPPSCs using Act A with or without other signaling factors to delineate the optimal conditional media sufficient to mediate DE- differentiation. Quantitative real-time PCR (qRT-PCR) analyses demonstrated a differential induction of the DE genes Forkhead Boxes A1 and A2 (FoxA1 and FoxA2) at day 5. FoxA1 transcripts elevated significantly in conditional media supplemented with Act A and Wnt3A, 14-fold relative to undifferentiated cells (Fig 1A). Similar induction levels were also observed in cells treated with Act A alone or Act A and BMP4, but albeit to a lower degree (6-to 8-fold) and notable variations between different experiments. Alternatively, FoxA2 expression levels were comparable in all tested conditional media, with a statistically significant 20-fold induction in media supplemented with Act A alone, and with Wnt3A or BMP4. Remarkably, conditional media supplemented with Act A and bFGF or FGF4 failed to significantly upregulate the studied DE-genes. Moreover, qRT-PCR analyses indicated an elevation in the transcript levels of GATA4, with a 40 to 60-fold increase relative to undifferentiated cells, post-treatment with Act A alone or in the presence of Wnt3A or BMP4 (Fig. 1A). In contrary, no GATA4 transcripts were detected in conditional media containing the three signaling factors together (Fig. 1A). Noticeably, Act A failed to induce GATA4 in media supplemented with either bFGF or FGF4 (Fig. 1A).

**Figure 1.**
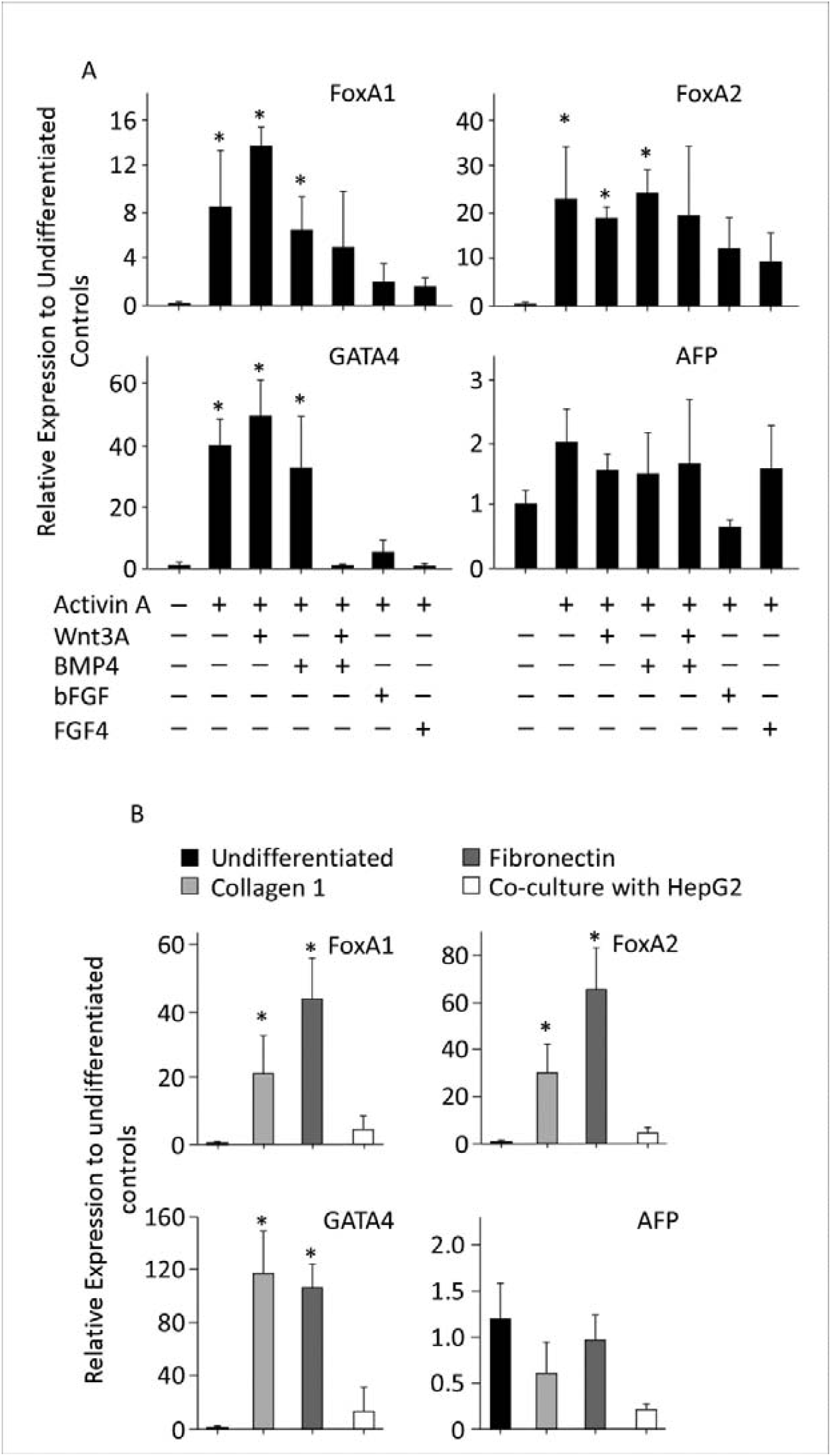
Molecular evaluation of different signaling factors and biomaterials on the generation of definitive endoderm lineage. qRT-PCR was performed for the indicated genes at day 5 of DPPSCs differentiation into DE lineage. A. Cells treated with differentiation conditional media supplemented with indicated signaling factors. B. Cells were seeded on precoated plates with/without the described scaffolds or co-cultured with HepG2 cells. Results were normalized to GAPDH and expressed relative to undifferentiated cells at day. Data are shown as mean ± SD (n = 4). *p < 0.05.

Worthy to note that at day 5, no significant upregulation of visceral endoderm and the early hepatogenic marker alpha-fetoprotein (AFP) in cells treated with the studied conditional media, suggesting that Act A and other signaling molecules do not support visceral endoderm lineage differentiation and are not sufficient for hepatogenic specification.

Taking together, the increase in expression levels of bona-fide DE- markers in conditional media supplemented with Act A alone or with Wnt3A suggests that our approaches using chemically defined-differentiation conditions has successfully enriched the DE- lineage of DPPSCs. Due to the experimental variations in the transcript levels of the DE- genes in cells treated with conditional media supplemented with containing Act A alone, therefore, the following experiments were carried out using Act A and WNT3A as the main induction cytokines.

It is well documented that of biomaterials in cell cultures supports viability, differentiation, maturation, and long-term function [36; 53]. To further improve the differentiation protocol and study the influence of biomaterials on DPPSCs mediated DE- indication, we perused DEdifferentiation under three different conditions: cells were cultured on plates precoated with collagen I, fibronectin, or co-cultured with HepG2 cells for 5 days.

Interestingly, the DE-marker gene expression was significantly elevated in DPPSCs differentiated on the scaffolds (Fig. 1B) versus on no scaffolds (Fig. 1A). Relative to DPPSCs differentiated on plain plates (Fig. 1A), FOXA1 transcript levels in cells differentiated on collagen I or fibronectin were 3- to 4-folds higher, respectively. Similarly, the expression levels of FOXA2 mRNA were elevated by 1.5- to 3-fold when cells were differentiated on the scaffolds (Fig. 1B). qRT-PCR analyses of GATA4 transcripts were significantly enriched in cells differentiated with collagen I or fibronectin, 120-fold increase relative to undifferentiation cells, at day 5 (Fig 1B). No DE-marker gene expressions were detected in DPPSCs co-cultured with HepG2, suggesting a differentiation suppression effect of hepatic cells (Fig. 1B). Furthermore, the used biomaterials did not facilitate prospective generation of visceral endoderm cells and did not support early hepatogenic specification, since AFP transcripts were not detected. In summary, biomaterial usage enhanced DPPSCs-mediated DE-differentiation. Since collagen I is an affordable, inexpensive substrate, and commonly used in stem cell studies, we use it as the scaffold to induce hepatogenic differentiation.

### 3.2. Hepatic Specification and Maturation

We investigated the developmental progress in the hepatogenic differentiation potential of DPPSCs using two-step protocol, which is schematically illustrated and presented in Fig. 2. At day 5 post DE-induction, the conditional media were supplemented with FGF4 and HGF, for 8 days, to enrich for hepatic cells specification, followed by 8 days culture in HCM supplemented with hepatogenic inducers such as OSM and Dex in addition to the growth factors (Fig. 2).

**Figure 2.**
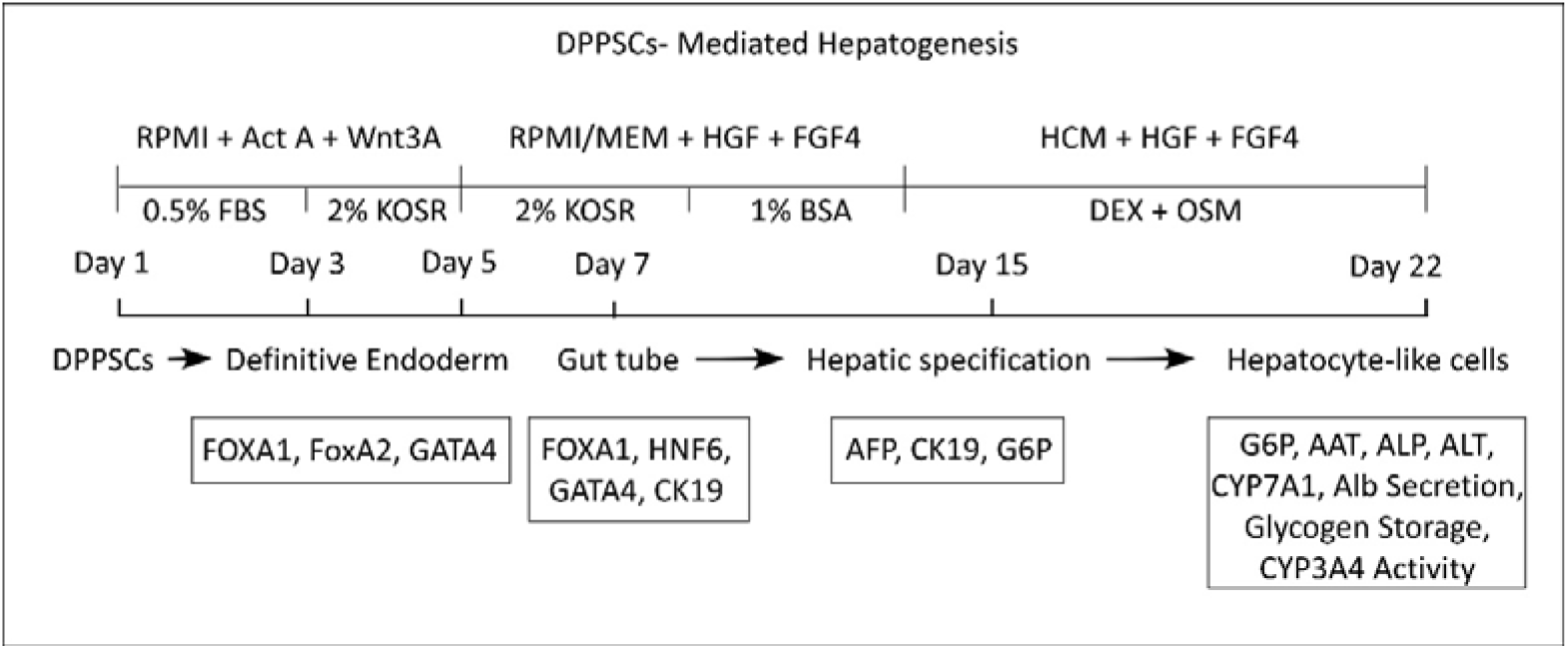
The experimental protocol for DPPSCs mediated- hepatocyte-like cell generation. Schematic representation of the differentiation protocol including the key manipulated signaling pathways and the different molecular and functional studies performed.

As we have previously reported, DPPSCs grow as a flat monolayer and exhibit a small spindle-shaped fibroblast-like morphology when cultured on polystyrene tissue culture plates or scaffolds (Fig. 3A) [25]. During the differentiation induction period, DPPSCs morphology further changed to a round or epithelioid shape, a characteristic of DE- cells Fig 3 B) [54]. At day 7, the cells became flat and tightly packed; and by day 22, they adopted a hepatocyte-like morphology (Fig 3 C-E). In contrast, undifferentiated cells cultured in regular growth medium containing FBS, leaned toward an elongated spindle-shaped fibroblastic morphology (Fig. 3F).

**Figure 3.**
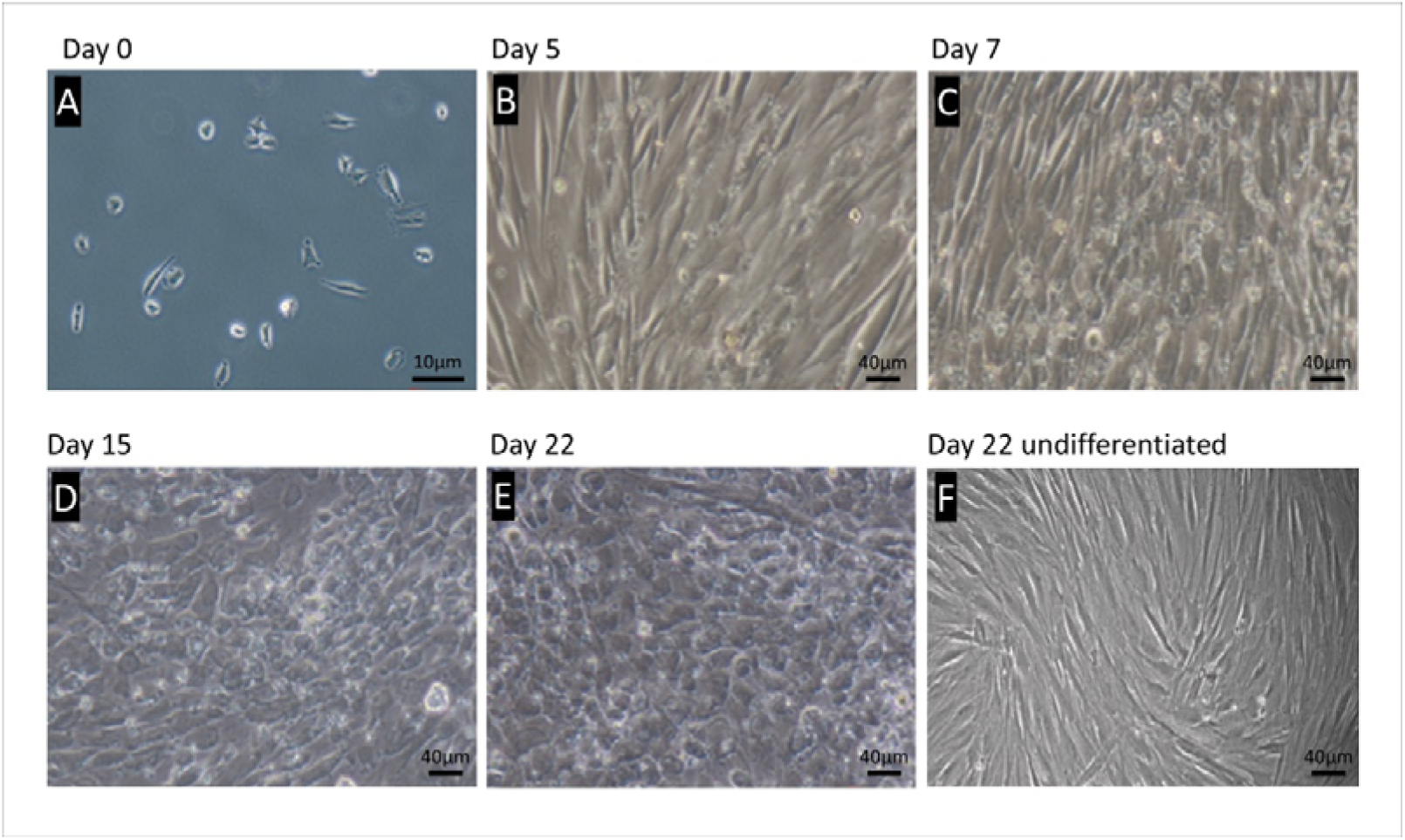
Changes of cellular morphological during the time course of DPPSCs- mediated hepatocyte-like. Phage contrast representative images of DPPSCs at different stages of differentiation from day 0 to day 22 and undifferentiated cells at day 22. Scale A. 100X magnification. B - F. 200X magnification.

At the molecular level, gene expression studies revealed a distinct early posterior foregut signature at day 7 of DPPSCs differentiation. The transcript levels of hepatic nuclear factor 6 (HNF6) and cytokeratin 19 (CK19) were notably elevated at day 7, relative to undifferenced cells (Fig. 4), indicating a significant commitment toward the foregut tube. Although the transcripts of GATA4 were upregulated at day 5 (Fig. 1B), simultaneously with the DE- markers, its elevated expression levels were maintained until day 7. Unlike, FoxA1 mRNA levels which were significantly reduced at day 7. This observation is likely due to a possible commitment of a small cell fraction to DE- lineage and/or a significant profile of these genes in the early posterior foregut (Fig. 4) [52; 55].

**Figure 4.**
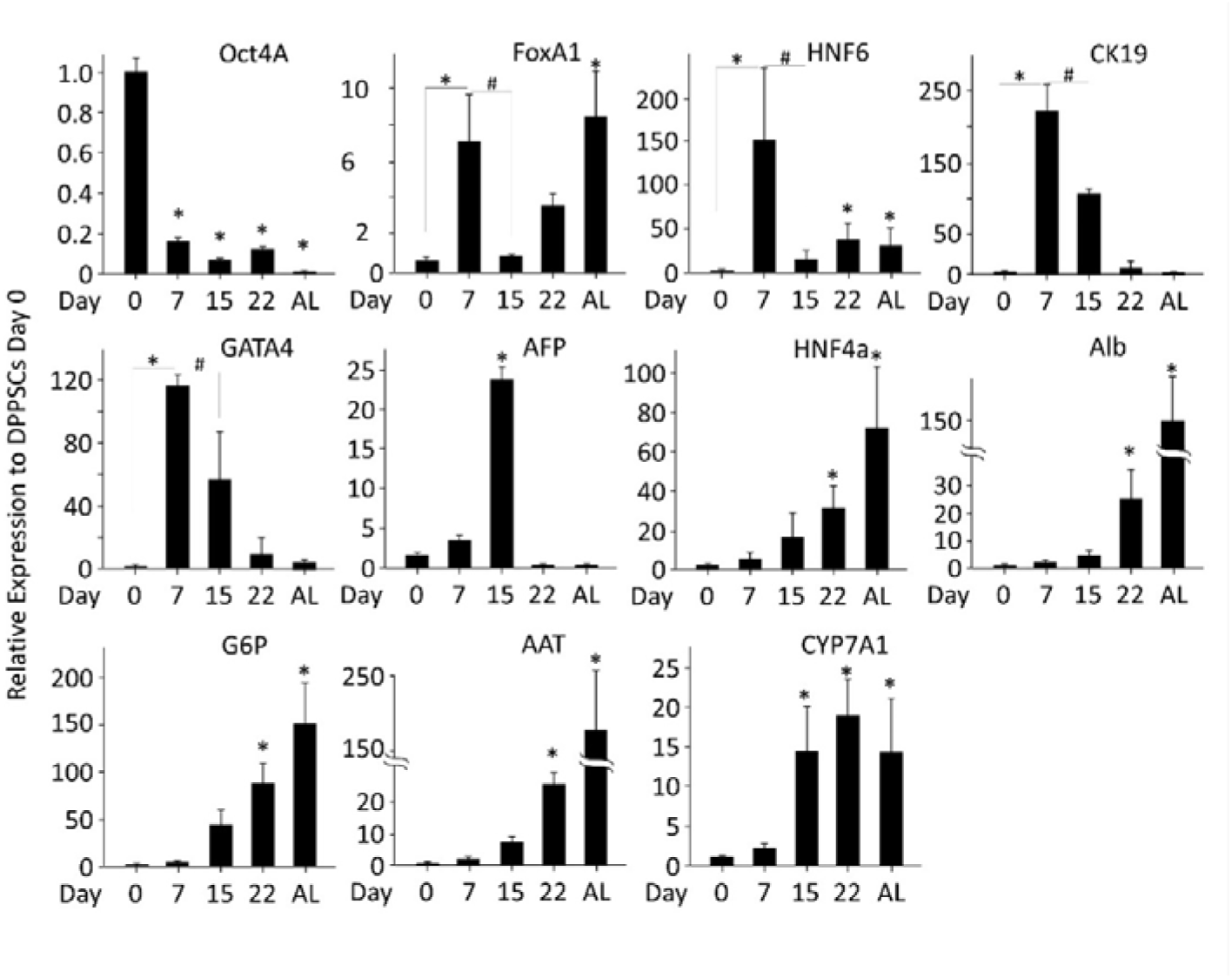
Molecular and cellular evaluation of DPPSCs- mediated hepatocyte-like cells. Time course qRT-PCR analysis for lineage-specific gene expression. The differentiation was performed using the previously described experimental protocol. Total RNA was harvested from the cells at the indicated days for gene expression. Results were normalized to GAPDH and expressed relative to DPPSCs at day 0, *p < 0.05. # p < 0.05 relative to Day 7. Data are shown as mean ± SD (n = 4).

Next, the developmental progress of differentiated DPPSCs toward hepatogenic specification was investigated at day 14 of treatment with conditional media. A notable reduction in the expression levels of the foregut tube markers was observed, concomitant with a significant increase in AFP transcripts (Fig. 4), which was not observed in earlier stages of differentiation, indicating a progression toward early hepatic specification. In addition, moderate levels of hepatic markers’ transcripts were noticed at day 14 (Fig. 4), including hepatic nuclear factor 4α (HNF4α), glucose 6-phosphate (G6P), and Cholesterol 7 alpha-hydroxylase (CYP7A1).

Simultaneously, eight days treatment of these committed DPSSCs with HCM supplemented hepatic induction molecules resulted in a significantly exaggeration in the expression levels of the hepatic markers at differentiation day 22. Relative to undifferentiated DPPSCs, the HNF4α and Alb transcripts were 40-and 15-fold higher, AAT and CYP7A1 transcripts were 20-fold higher, and G6P mRNA levels were 100-fold higher at day 22 (Fig. 4). Taking together, we implemented an optimized 3-stage protocol that directed a stepwise differentiation of DPPSCs to the hepatocyte-like cell populations.

### 3.3. DPPSCs-derived hepatocyte-like cells demonstrate hepatic function

Next, to corroborate the qRT-PCR analysis and evaluate the differentiation efficiency, we ascertained the protein expression of the hepatogenic markers by immuno-localization and functional analysis. Immunofluorescence with specific antibodies directed against AAT and Alb proteins revealed a significant expression of these hepatic markers at day 22 of DPPSCs differentiation (Fig. 5A-B).

**Figure 5.**
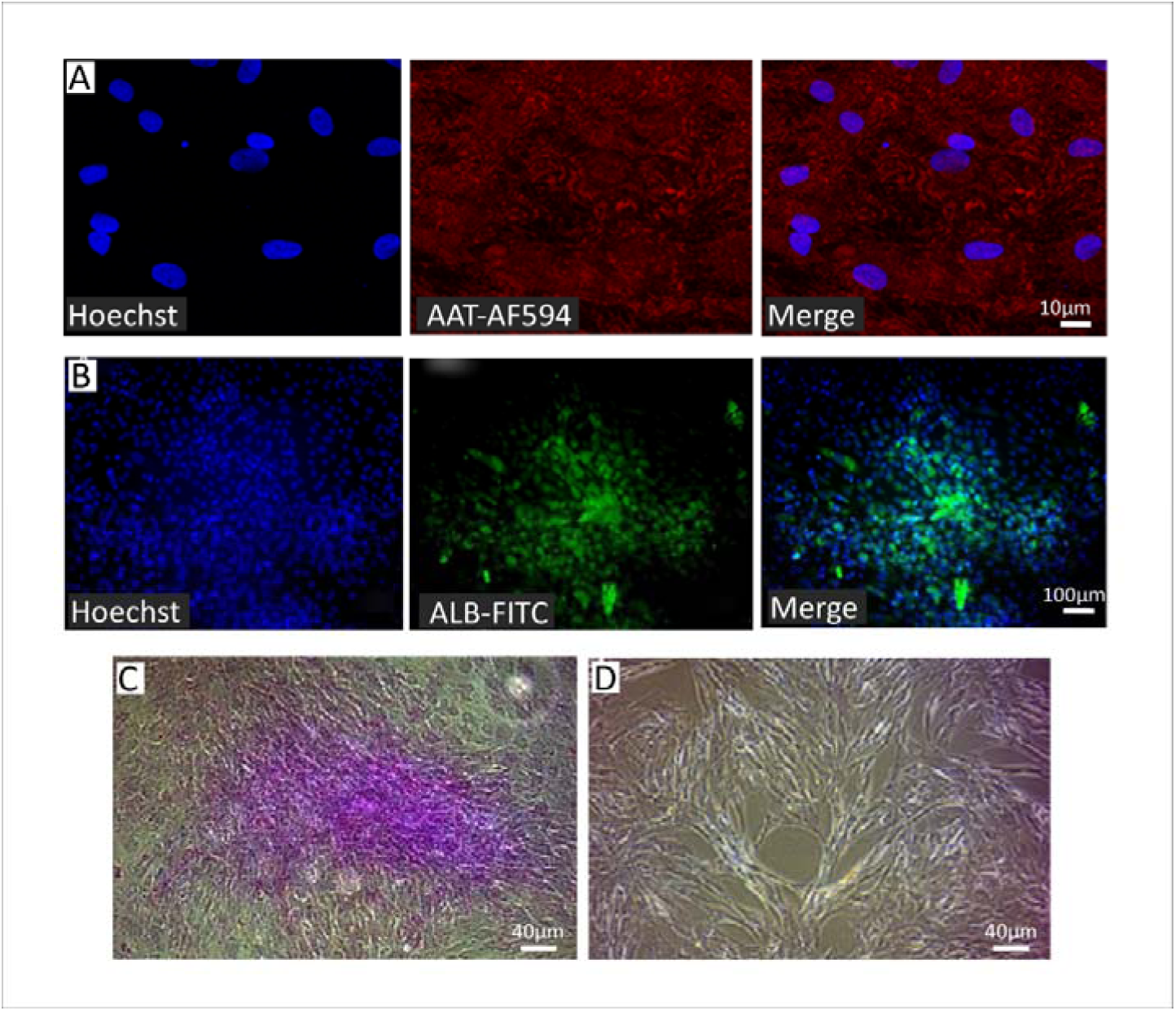
Protein expression and glycogen storage of DPPSCs- mediated hepatocyte-like cells. A. and B. Representative fluorescence microscopy images of DPPSCS mediated hepatocyte-like cells at day 22, immunoassayed with antibodies directed against A. AAT (AF594, red) and B. ALB (FITC, green) proteins. Hoechst nuclear marker in blue. 200X magnification. C. and D. Representative light microscope images of differentiated and undifferentiated DPPSCs at day 22, respectively, stained with periodic acid Schiff (PAS) for glycogen storage detection. 100X magnification.

Mature hepatocytes are characterized by their ability to store glycogen [56]. Interestingly, most of the differentiated DPPSCs showed a significant positive periodic acid Schiff (PAS) staining in the cytoplasm, albeit to a different degree, at Day 22 (Fig. 5C), which is indicative for glycogen storage, and the generation of mature hepatocyte-like cells. Alternatively, no PAS staining was detected in undifferentiated cells (Fig. 5D).

Then, we investigated the activity of hepatic metabolic enzymes. Time course studies performed on protein extracts from differentiated DPPSCs for GGT, AST, ALP, and ALT activities revealed a time-dependent upregulation of liver metabolic enzymes which beaked at day 18 of differentiation (Fig. 6A-D, respectively). Furthermore, the enzymatic activity of CYP3A4 was significantly induced in differentiated DPPSCs post treatment with rifampicin, 5-fold relative to untreated cells. Co-treatment with erythromycin, a CYP3A4 inhibitor, significantly down-regulated the observed enzymatic action against rifampicin (Fig. 6E).

**Figure 6.**
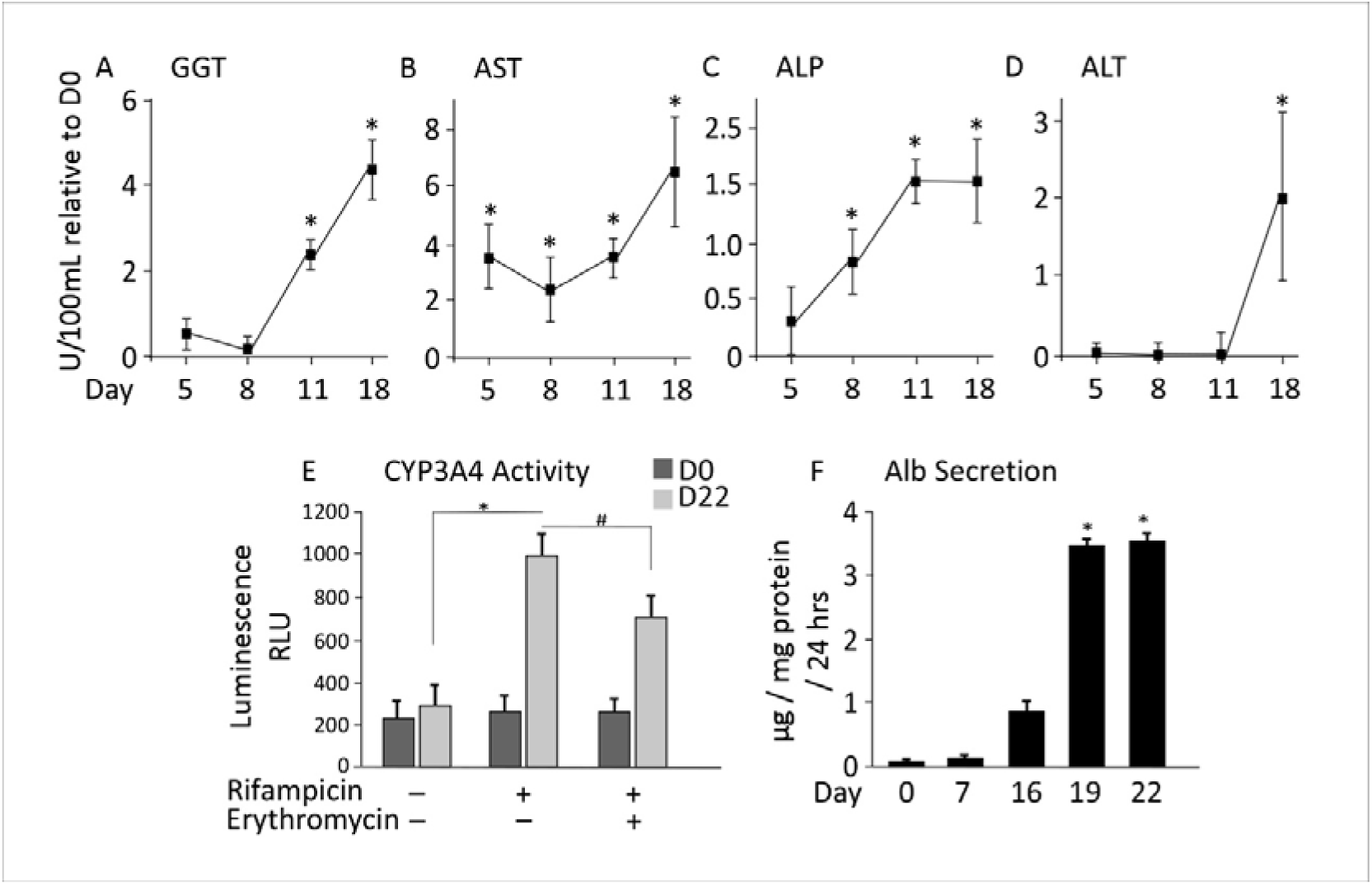
Functional analysis of the generated DPPSCs- mediated hepatocyte-like cells. Time course hepatocyte enzymatic activity of A. GGT, B. AST, C. APF, D. ALT were performed for the differentiated DPPSCs. A notable induction was observed at day 18. E. CYP3A4 activity was measured by luminescence in DPPSCs at day 0 and day 22 of differentiation using rifampicin in the presence or absence of the antagonist erythromycin. F. Alb secretion assay was performed at a time course differentiated DPPSCs, elevated level of secreted Alb was detected at day 19 and later. All measurements are relative to DPPSCs at Day 0, * p<0.05. # p < 0.05 relative to rifampicin treatment alone. Data are shown as mean ± SD (n = 4).

Alb secretion was assayed at the time course of the differentiation process. As shown in Fig. 6F, at earlier stages of hepatogenesis, day 16 of differentiation, a limited amount of secreted Alb was detected in the media. However, at differentiation day 19 and 22, the amount of secreted albumin was triplicated into 3.5 ug/mg protein (Fig. 6F). Taking these observations together; the described differentiation protocol (Fig. 2) efficiently induced the generation of DPPSCs mediated functional hepatocyte-like cells and indicates that DPPSCs are a useful cell models for hepatogenesis.

## 4. Discussion

The current strategy in stem cell differentiation protocols is to mimic cellular signaling events associated with the embryonic developmental process for the lineage of interest [8, 47, 48]. However, *in vitro* cell differentiation gives a broad range of outcome lineages, thus, the development of a protocol that significantly improves a targeted cell type is of particular interest [49]. Targeted differentiation requires culturing the cells in chemically well-defined media, which regulate cell signaling pathways, resulting in the generation of cell type of interest. This approach has been successfully applied to hESCs and iPSCs [51–53]; but has yet to be applied to other pluripotent cell types, including adult DPPSCs.

In this study, we applied an optimized 3D-3-stages protocol directed toward a stepwise differentiation of DPPSCs to hepatocyte-like cells. For efficient differentiation, cells were checked at each developmental stage using lineage-specific markers. Initially, DPPSCs were cultured for 5 days in differentiation media supplemented with Act A, with or without Wnt3a, BMP4, or the growth factors bFGF and FGF4. Interestingly, Act A alone was sufficient to induce DE-differentiation. Our results are in accordance with the reported role of Act A inducing DE from pluripotent stem cells [55; 57]. It is well known that nodal signaling is required for endoderm specification during the gastrulation [58]. Furthermore, DE-induction depends on the duration of exposure to the stimulation factors; Agarwal *et. al*. reported that 5-days incubations are sufficient to induce DE [49].

Media supplemented with Act A and Wnt3A were superior in inducing DE-lineage [57; 59]. Using hESCs, Mathew *et. al*. observed an enhanced DE induction using Act A and Wnt3A supplemented media compared to media contained Act A alone [51]. Like hESC, in this study, we observed an improved induction of DE-markers in DPPSCs cultured with Act A and Wnt3A. Alternatively, the growth factors bFGF or FGF4 suppressed Act A-mediated DE induction [60]. Noteworthy, FGF family members are multifunctional factors. They can maintain hESC pluripotency [61] or DE-mediator [62], depending on the associated conditions.

During liver development, BMPs signaling is crucial; however, their functional role during *in vitro*-adult stem cell differentiation is not clear. Cultural media supplemented with BMP2 and BMP4 fail to improve hepatogenesis from adipose tissue stem cell [63]. In accordance with the other studies, our results confirmed BMP4 signaling has no functional role during DPPSCs differentiation into DE reconciled by Act A induction. Otherwise, studies have shown that a combination of Act A and BMP4 signaling induce mesoderm formation [64]. Notably, DE induction was further enhanced using three-dimensional (3D) scaffolds. Treatment with either collagen or fibronectin upregulated DE markers indicated that 3D enhances the cellular programming potential, which is in accordance with previously reported observations using ESCs, iPSCs and other mesenchymal stem cells [36; 37; 65].

The use of minimal amount of FBS in differentiation media has been previously reported to inhibit PI3K signaling [66], which is necessary to enhance ACT A-mediated DE generation. Using 0.5% FBS during the first 5 days of DE induction, D’Amour and her team observed an enhancement in the expression of Sox17 or FoxA2 in hESC [57]. In our protocol, we used 0.5% FBS-media for the first 3 days of differentiation. Then, it was replaced with 2% KOSR for an extra 4 days, a serum that has been proven to be effective and suitable with defined components [49].At differentiation days 7-22, KOSR was removed and hepatogenic lineage induction was carried on with serum-free conditional culture media. As descripted in schematic Fig. 2, the growth factors FGF4 and HGF were used to induce hepatogenesis, followed by the usage of OSM with Dex for the hepatocytes’ maturation. The role of FGF4 and HGF in hepatocyte specification has been previously reported for both Wharton’s jelly- and bone marrow mesenchymal stem cells [67; 68; 69]. In accordance, both FGF4 and HGF were required to mediate hepatic specification in DPPSCs. We observed an improved expression of several hepatic markers including immature and mature markers at day 13 of differentiation.

Studies have shown that the interleukine-6 family cytokine (OSM) is required for hepatocyte maturation in combination with glucocorticoids, such as Dex [70; 71]. Additionally, it is well documented that HGF, EGF, and OSM have decisive effects on the maintenance of primary human hepatocytes *in vitro* [72]. The combination of HGF, OSM, and Dex is widely used in protocols to differentiate stem cells into hepatocytes [49; 73]. In our protocol, we applied the maturation factors OSM and Dex at day 13 posting commitment to the hepatic fate. Consequently, we documented hepatocyte-like cell maturation using cellular and molecular techniques. After 15 days exposure to media containing OSM and Dex, we observed changes in cell morphology with hepatocyte-like structures, which was associated with Alb secretion, and the activation of hepatic specific enzymes.

After the establishment of the most optimal protocol used for DPPSC, we performed the tests that were available for us to prove the efficacy of the differentiation of DPPSC into hepatocyte-like cells. At the molecular level, differentiated DPPSC expressed AAT, Alb, and G6P transcripts, genes that are markers for mature hepatocytes. Moreover, the protein expression of these markers was also documented in our study. Cellular morphological changes were also detected, and mature cells acquired polygonal-shape typical for hepatocytes [74]. In addition, DPPSCs mediated hepatocyte-like cells were functional. These cells stored glycogen, secreted Alb, and other metabolic enzymes as efficient as other stem cell models used for hepatogenesis including MSCs [39] and hESCs[75; 76]. Furthermore, differentiated DPPSCs have shown an improved Cyp3A4 enzymatic activity, suggesting that the generated hepato-cyte-like cells can be used as a model for drug metabolism, which would be useful for the pharmaceutical industry [77].

## 5. Conclusions

We designed a novel protocol to differentiate DPPSCs into functional hepatocyte-like cells. Our protocol is a directed differentiation protocol mimicking the stepwise process observed during embryonic liver development. Using DPPSCs as a pluripotent-like stem cell model suggesting their potential usage for liver regenerative medicine and prospective treatments to be developed in the future. However, for clinical applications, there are still many studies that are needed to be improved, yet the generated functional DPPSCs-hepatocyte-like cells can be used as a model for drug screening, hepatic metabolism studies, and hepatic disease applications.

## Data Availability

All data are available.

## Acknowledgment

Authors would like to thank Dr. Cámara Vallejo at Oral and Maxillofacial Surgery Department, Hospital Clinico de Barcelona, Barcelona, Spain, for patient’s recruitment and molar extractions.

## Abbreviations

AST: Aspartate aminotransferase
BAL: Bioartificial liver
CGH: Chromosome genomic
hybridization: 
DEX: Dexamethasone
DPMSC: Dental pulp mesenchymal stem cells
DPPSC: Dental pulp pluripotent stem cells
EB: Embryoid bodies
ECM: Extracellular matrix
ESDL: End-stage liver disease
GGT: Gammaglutamyl transferase
HB: Hepatoblasts
HCC: Hepatocellular carcinoma
HT: Hepatocyte transplantation
MSC: Mesenchymal stromal cells
OLT: Orthotopic liver transplantation
OSM: Oncostatin M

## Authors’ contribution

C.G.R., Collection and assembly of data and first manuscript draft; S.M. and S.A.D., Data analysis and interpretation; M.A.A. and C.M., Statistic analysis; E.F.P. and E.F.A, Provision of study patients and cells isolation; M.B., A.A.M and M.A., Conception and design, collection and assembly of data, data analysis and Interpretation, manuscript writing, and final approval of the Manuscript.

